# Maintaining and escaping feedback control in hierarchically organised tissue: a case study of the intestinal epithelium

**DOI:** 10.1101/2021.06.11.448040

**Authors:** Matthias M. Fischer, Hanspeter Herzel, Nils Blüthgen

**Affiliations:** Institute for Theoretical Biology, Charité and Humboldt Universität zu Berlin, 10115 Berlin, Germany

**Keywords:** Cancer, Colon, Dedifferentiation, Intestinal epithelium, Regeneration, Tissue homoeostasis

## Abstract

The intestinal epithelium is one of the fastest renewing tissues in mammals. It shows a hierarchical organisation, where intestinal stem cells at the base of crypts give rise to rapidly dividing transit amplifying cells that in turn renew the pool of short-lived differentiated cells. Upon injury and stem-cell loss, cells can also dedifferentiate. Tissue homeostasis require a tightly regulated balance of differentiation and stem cell proliferation, and failure can lead to tissue extinction or to unbounded growth and cancerous lesions. Here, we present a two-compartment mathematical model of intestinal epithelium population dynamics that includes a known feedback inhibition of stem cell differentiation by differentiated cells. The model shows that feedback regulation stabilises the number of differentiated cells as these become invariant to changes in their apoptosis rate. Stability of the system is largely independent of feedback strength and shape, but specific thresholds exist after unbounded growth occurs. When dedifferentiation is added to the model, we find that the system can recover more gracefully after certain external perturbations. However, dedifferentiation makes the system more prone to loosing homoeostasis. Taken together, our mathematical model shows how a feedback-controlled hierarchical tissue can maintain homeostasis and can be robust to many external perturbations.

## I. INTRODUCTION

In a hierarchically organised tissue, usually two classes of cells can be distinguished: Adult stem cells with unlimited capacity of self-renewal, which differentiate into cells with limited proliferative potential that perform the designated function of the tissue [1]. Additionally, tissue damage can lead to dedifferentiation of differentiated cells back into cycling stem cells Bonventre [2], Tata et al. [3], Liu and Chen [4], Liu et al. [5]. Homoeostasis of such a tissue in the face of external perturbations requires a tight regulation of the stem cell compartment. Upon tissue damage, stem cells need to increase proliferation; however proliferation of the stem cell compartment has to be tightly controlled to prevent unlimited growth of the tissue [6]. Such a control seems to be realised by specific feedback loops exerted by differentiated cells onto the stem cell compartment [7]. In contrast, control of the dedifferentiation of differentiated cells seems to be exerted by the stem cell compartment (Tata et al. [3], Beumer and Clevers [8] and the references therein).

The intestinal epithelium is a prime examples of hierarchically organised tissues. Despite its simple single-layered structure, it is able to withstand continuous mechanical, chemical and biological stresses due to its specific tissue architecture and a high rate of tissue turnover [9]: Stem cells at the bottom of the intestinal crypts divide continuously approximately once per day. Cells mature while migrating out of the crypt until they terminally differentiate and become part of the villi, eventually undergoing apoptosis and being shed into the intestinal lumen [10]. In the intestine and colon, the stem cell compartment is controlled via differentiated epithelial cells releasing Indian Hedgehog (Ihh), which stimulates mesenchymal cells to release Bone Morphogentic Proteins (BMPs). These interfere with intracellular effects of WNT signalling and thus stimulate stem cell differentiation [11–15].

Previous theoretical research on hierarchical tissues has often focussed on the case of arbitrary tissues: Rodriguez-Brenes et al. [16] considered a generic hierarchival tissue with a compartment of cycling and differentiating stem cells, and another compartment of non-cycling differentiated cells undergoing apoptosis, assuming the differentiated cell compartment may exert feedbacks onto the stem cells by decreasing their rate of proliferation and reducing the probability of symmetrical stem cell division. They then studied the order in which mutations in feedbacks need to arise to yield a selective advantage. The same model was used in Rodriguez-Brenes et al. [17] to reveal that during recovery from an injury dampened oscillations may occur, which is more pronounced when the stem cells are only a small fraction of the cell population. The same model has also been studied by Sun and Komarova [18] to obtain analytical solutions for the mean and variance of the cell compartment sizes. Recently, Wodarz [19] has extended the model by including dedifferentiation. Assuming a specific sigmoidal feedback onto stem cell cycling rate and self-renewal probability, he studied the effect of a linear and a sigmoidal dedifferentiation term, showing how unbounded cancerous tissue growth may arise as a consequence of escaping this feedback. He also demonstrated how dedifferentiation may allow for speedier regeneration after perturbations using numerical simulations.

Other tissue-specific theoretical studies include models of the haematopoietic system [20–24], the mammalian olfactory epithelium [25, 26], and breast cancer [27]. Models of the intestinal epithelium have been reviewed in Carulli et al. [28]. These include Paulus et al. [29], a model for its post-irradiation recovery that showed that damage-control response may reside solely in the stem cell compartment without requiring a feedback from the villus, challenging earlier views. Another important model of the colon epithelium [30] showed that a ”bang-bang” control of proliferation is optimal to minimise the time for crypt construction. However, this model assumes continuous exponential growth, thus it is not suitable to address tissue homoeostasis and stability.

Theoretical examinations that address mechanisms of homoeostasis and dynamics of healthy intestinal epithelium have been rare. A noteworthy exception is Johnston et al. [31], who derived a three-compartment ODE model distinguishing stem, transit-amplifying and terminally differentiated cells, assuming that the first two compartments limit their own growth via a negative saturating or a negative quadratic term, reminiscent of classical single-species population dynamic models [32]. However, no mechanistic justification of these assumptions has been provided, possibly owing to the fact that our mechanistic understanding of the biology of intestinal stem cells has been limited until very recently [9]. Therefore, in this work, we set out to close this gap in the literature and derive a model of intestinal epithelial population dynamics based on our current mechanistic understanding of the involved underlying biological processes. This model allowed us to study specifically the role of feedbacks and dedifferentiation in tissue homoeostasis and regeneration for the specific case of the colon epithelium.

## II. MATERIALS AND METHODS

### A. Colon epithelium model

Our main model consists of two cell compartments *S* and *D*, denoting stem cells, and differentiated cells, respectively, following earlier approaches Rodriguez-Brenes et al. [16]. Stem cells cycle at a constant rate *β*, differentiated cells are cell-cycle arrested and die with an apoptosis rate *ω*. Finally, stem cells differentiate with a rate *δ*(*D*) that is a function of the size of the differentiated cell compartment. Following earlier models in the literature such as Rodriguez-Brenes et al. [16], we consider transit-amplifying cells to be a part of the differentiated cell compartment *D*, since their limited proliferative potential distinguishes them from stem cells [9].

Overall, the model is described by two coupled ordinary differential equations:

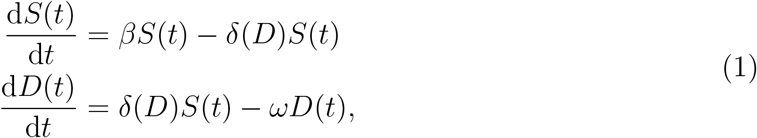

where 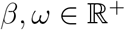 and *δ* is a continuously differentiable, monotonically increasing, invertible function 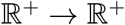. This function *δ* models the results of experimental studies on intestinal tissue that demonstrated the stimulation of stem cell differentiation by differentiated cells via Indian Hedgehog (*Ihh*), and Bone Morphogenic Protein (*BMP*) [11, 12]. A sketch of the model is shown in Figure 1a..

**FIG. 1.**
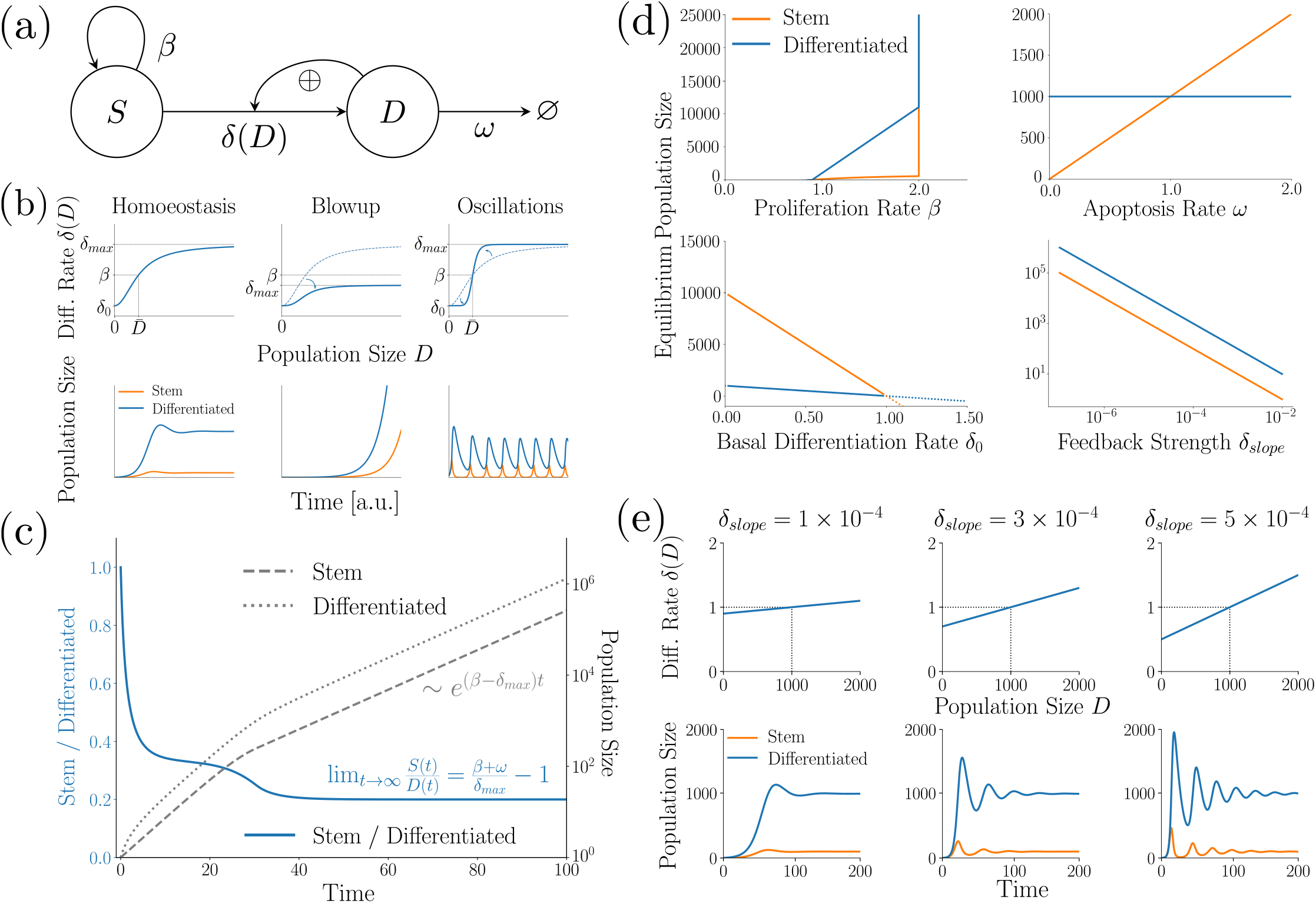
The basic colon epithelium model. **(a)** Schematic sketch of the model. **(b)** The feedback function *δ* (upper row) determines the qualitative behaviour of the model (lower row). First column: Stable steady-state. Second column: unbounded growth. Third column: sustained oscillations. **(c)** In case of explosive growth, the system converges to a stable ratio of stem to differentiated cells. Exemplary numerical integration. **(d)** Bifurcation diagram of the system with linear feedback function *δ*. Standard parametrisation: *β* = 1.0, *ω* = 0.1, *δ*_0_ = 0.9, *δ_slope_* = 10^−4^, *δ_max_* = 2.0. Note the switch from a stable steady-state to unbounded growth for *β* > *δ_max_* (upper-left panel). Also note the invariance of 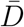 to changes in *ω* (upper-right panel). Both properties hold regardless of how we chose *δ*. Finally, notice that *δ*_0_ *>* 0, *δ_slope_* > 0 can be made arbitrarily small without switching to unbounded growth; however, if *δ*_0_ > *β*, the system will go extinct (lower two panels). **(e)** A steeper feedback functions causes stronger and longer oscillations. Exemplary numerical simulations with linear function *δ* for varying *δ_slope_*; steady-state fixed at 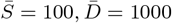.

### B. Colon epithelium model with dedifferentiation

We extend the model by a dedifferentiation process of differentiated cells into stem cells. In this model, the rate of dedifferentiation is a continuously differentiable and monotonically decreasing function 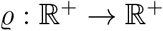 of the stem cell compartment, where a higher number of stem cells reduces dedifferentiation. Thus, the model reads:

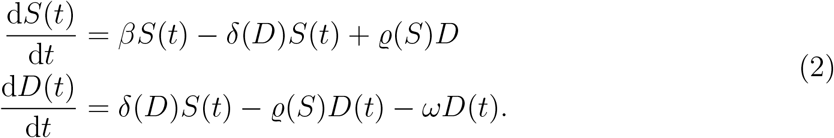

A sketch of the model is shown in Figure 4a.

### C. Numerical simulations

Numerical simulations have been implemented in the Python programming language, version 3.7.3. We provide the complete commented source code under: https://github.com/Matthias-M-Fischer/Epithelium.

## III. RESULTS

### A. A population dynamics model of the colon epithelium

We developed a model of the intestinal epithelial population dynamics (Fig. 1a, ‘Materials and Methods’), distinguishing between two compartments: Stem cells cycling at a rate *β* and differentiating at a rate *δ*(*D*); and differentiated cells undergoing apoptosis and leaving the system at a rate *ω*. We model the differentiation rate as a positive function of the size of the differentiated cell compartment *D*, reflecting earlier experimental results [11, 12].

#### 1. Stability can be lost via two routes

First we study the steady-states of system (1). For now, we will not use any specific function *δ* and only demand that *δ* is a positive, monotonic, continuously differentiable and invertible function. We also assume d*δ*/d*D* ≥ 0, since differentiated cells stimulate stem cell differentiation.

By solving d*S*(*t*)/d*t* = d*D*(*t*)/d*t* = 0 we find two steady-states, which in the following we will mark with an over-bar. We find a trivial steady-state 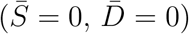 that exists for all parameter values. Under some conditions, also a non-trivial steady-state exists at 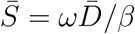 and 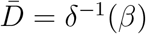. Here, *δ*^−1^ denotes the inverse function of *δ*. This non-trivial steady-state exists if *δ* can reach the value of *β*. Thus, this second steady-state is only present if the maximum differentiation rate is higher than the proliferation rate. One may see that the steady-state does not depend on the apoptosis rate *ω*. This is a biologically interesting property which we will return to later.

To address the stability of the steady-states, we investigate the Jacobian [33] of the system:

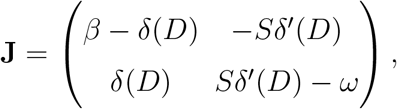

which at the trivial steady-state has eigenvalues λ_1_ = *β* – *δ*(0), λ_2_ = −*ω* < 0. Hence, the trivial steady-state is only stable if *β* is smaller than the basal differentiation rate *δ*(0).

At any non-trivial steady-state, the Jacobian is given by

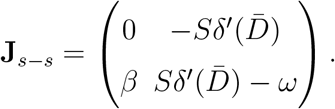

The steady-state is stable if det **J**_*s–s*_ > 0 and tr **J**_*s–s*_ < 0 (Routh–Hurwitz criterion, see Strogatz [33]), which leads to the condition that 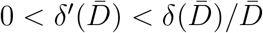.

These results reveal two routes of how the intestinal epithelium can lose homeostasis (Fig. 1b): First, the tissue might show unbounded growth, which could be interpreted as the emergence of a cancerous lesion (Fig. 1b, second column). This occurs once the proliferation rate *β* exceeds the maximum differentiation rate *δ_max_* (either by increased proliferation or decreased maximal differentiation). In this case, only the (unstable) steady-state (0, 0) remains, and any positive perturbation of *S* away from it will lead to unbounded growth.

Second, the steady-state 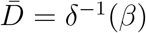 can get unstable if 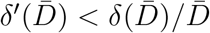. In other words, at the steady-state the slope of the feedback function might exceed 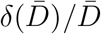. Then, the behaviour of the system depends on the feedback function. This can be illustrated with the following two examples:

First, consider a linear function of the form *δ*(*D*) = *δ*_0_ + *δ_slope_D*. The intercept *δ*_0_ denotes the basal differentiation rate of stem cells in the absence of external stimuli, whereas *δ_slope_* quantifies the increase in differentiation as a function of *D*(*t*). For instability, we require 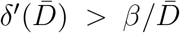, hence the system will only be stable for a strictly positive *δ*_0_. This is biologically plausible, since the basal stem cell differentiation rate can be expected to exceed zero. If, however, *δ*_0_ = 0, then the fix point will be unstable. This case is equivalent to the classical Lotka-Volterra model of predator-prey population dynamics, exhibiting undampened oscillations. As a second example, consider a sigmoidal function of the form 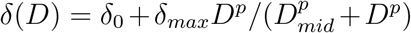. Here, *δ_max_* denotes the maximum increase in differentiation rate the stem cell compartment may experience, *D_mid_* is the position of the mid-point of the sigmoidal function, and *p* quantifies the steepness of the feedback function around it. For instability, we require 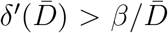. Hence, at *p* > 4*β*/*δ_max_* the steady-state loses its stability, and sustained oscillations start to occur.

#### 2. Exponential growth and convergence to a stable cell type ratio after escaping control

Next, we analyse the behaviour of system (1) during unbounded growth. This case corresponds to a tissue that has escaped homeostatic control and has degenerated into a cancerous lesion. In our model, this occurs when *β* exceeds the maximum of *δ*, implying that *δ* has a maximum value *δ_max_* < *β*. Such saturation of the differentiation rate could arise from saturated signalling or thermodynamical constraints.

Interestingly, during such unbounded growth the system will converge to a stable cell type ratio (*S*(*t*)/*D*(*t*) = const for *t* → ∞): After feedback saturation, the dynamics of the system is governed by the following differential equations:

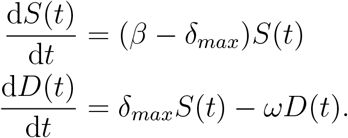

Directly solving for *S*(*t*) yields a simple exponential function, and subsequently we may also solve *D*(*t*). Overall, we get:

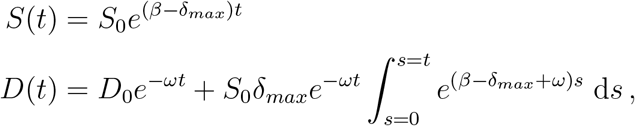

where *S*_0_ = *S*(0), *D*_0_ = *D*(0) denote the initial condition of the system. This yields

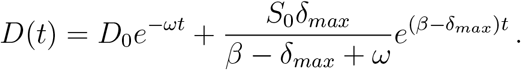

This allows us to take the following limit

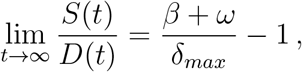

which denotes the ratio of stem to differentiated cells the system will converge to during explosive growth.

Additionally, one may see that after the transient period the overall growth of the system amounts to

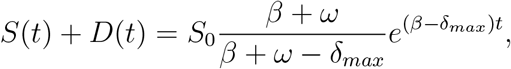

which implies an exponential growth at a constant rate *β* – *δ_max_*. Figure 1c provides an exemplary numerical simulation of the system with a piecewise linear feedback function *δ*(*D*) = min{0.9 + 10^−4^*D*, 1.0}, and parameters *β* = 1.1 and *ω* = 0.1. Observe the convergence of the ratio *S*(*t*)/*D*(*t*) to a stable ratio of −1 + (*β* + *ω*)/*δ_max_* ≈ 0.2, which is in agreement with our theoretical analysis.

#### 3. Bifurcation analysis for the case of piecewise linear δ

Now, we further analyse the influence of system parameters on the behaviour of system 1. To this end, we now choose a concrete differentiation rate function *δ*. For simplicity, we choose a piecewise linear function with a positive intercept *δ*_0_ > 0, denoting the basal stem cell differentiation rate. We assume that *δ* grows linearly in *D* with a slope of *δ_slope_* > 0, until it reaches an upper bound *δ_max_*, where it stops increasing. Hence, we have

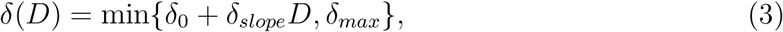

where *δ*_0_, *δ_slope_*, *δ_max_* > 0.

The non-trivial steady-state resides at 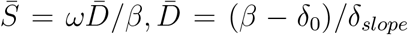 and only exists if *β* < *δ_max_*. The steady-state is only biologically feasible 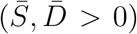 if *δ*_0_ < *β*. Then, the Jacobian at the steady-state is given by

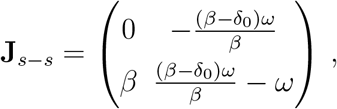

with eigenvalues

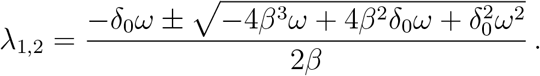

If

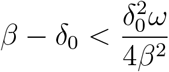

the radicand is positive. Then, due to *δ*_0_ < *β* both eigenvalues will be negative and real, hence the steady-state is a stable node. If, however, the radicand is negative, oscillations occur and the real part of the eigenvalues is always negative due to *δ*_0_, *ω* > 0, making the steady-state a stable focus. In any case, any biologically feasible steady-state of this system will always be stable.

Biologically, it is interesting that no other constraints on *δ*_0_ and *δ_slope_* are required except for *δ*_0_ < *β*, which we require for a feasible steady-state, and *β* < *δ_max_*. In particular, *δ*_0_ > 0, *δ_slope_* > 0 can be arbitrarily small, yet the system remains stable. Similarly, changes in *ω* ≥ 0 will not affect the stability of the system. Figure 1d illustrates these findings and shows bifurcation diagrams of the system for varying parameters *β*, *ω*, *δ*_0_, *δ_slope_*.

#### 4. The feedback loop can cause oscillatory behaviour in cell numbers

Next, we investigate the case of the system showing dampened oscillations around the steady-state. For any given steady-state 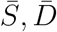 it holds that 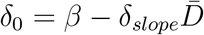. This permits to express λ_1,2_, and thus the amplitude decay and the angular frequency of the dampened oscillations in terms of *δ_slope_*. We find that the amplitude of the oscillations will decay with

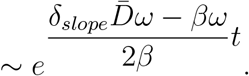

Note that at any equilibrium the exponent is negative because from 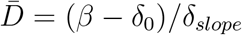 it follows that 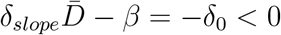. The angular frequency of oscillations reads

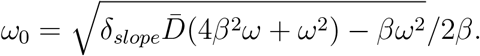

Hence, a greater slope of the feedback function *δ* will result in a slower decay of the amplitude, as well a higher cycling frequency. For illustration, we provide a set of numerical simulations of the system with a piecewise linear function *δ* as defined before and varying values of *δ_slope_* in Figure 1e.

The relationship between function slope and oscillatory behaviour of the system is a biologically interesting result. Observe how large values of *δ_slope_* cause strong oscillations which are biologically unlikely. This suggests that to minimise oscillations in cell numbers, a smaller slope of the feedback function is desirable. For very small slopes, the difference between *β* and *δ*_0_ needs to become sufficiently small as well, if the position of the steady-state 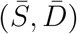 should remain constant. Then, the oscillations will vanish, and the steady-state becomes a stable node, as shown previously.

### B. Comparison of different model topologies

In the previous section, we have shown that the colon epithelium model can cause homoeostasis through a feedback from differentiated cells onto the stem cell differentiation rate. Now, we explore how alternative feedback topologies in our two-compartment colon model might alter tissue homeostasis, and compare the properties of these models. We therefore generate the family of all six one-looped model topologies.

#### 1. Controlling stem cells is required for stability

First, we consider those two model topologies where the apoptosis rate *ω* is regulated by either *S* or *D*, respectively. These feedback loops are biologically unlikely, but we mention them for the sake of completeness. Both topologies are not able to show homoeostasis: If both *β* and *δ* are unregulated and hence constant, we have d*S*(*t*)/d*t* = *βS* – *δS*, which for any non-trivial steady-state requires *β* = *δ*. This, in turn, implies ∀*t* : d*S*(*t*)/d*t* = 0, hence after any perturbation of *S, S* will not return to its pre-perturbation state. Accordingly, these models are biologically not realistic, and we will not consider them in the rest of this paper.

#### 2. Indirect regulation of the stem cell compartment decouples the steady-state number of differentiated cells from their apoptosis rate

Next, we show that all models in which the stem cell compartment is exclusively regulated by the differentiated cell compartment *D* following arbitrary continuously differentiable functions *β*(*D*), *δ*(*D*) enjoy the property of 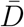 being independent of *ω*: Let d*S*(*t*)/d*t* = *β*(*D*)*S* – *δ*(*D*)*S*. At any non-trivial steady-state 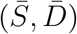, from d*S*(*t*)/d*t* = 0 we get that 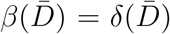. Define *α*(*D*) := *β*(*D*) – *δ*(*D*), then 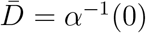, which does indeed not depend on our choice of *ω*.

#### 3. Saturating feedback functions β and δ cause convergence to a stable cell type ratio during unbounded growth

Finally, it may also be briefly remarked that all systems in which proliferation and differentiation rates of the stem cell compartment are functions saturating to *β_min_* and *δ_max_* will in case of unbounded growth converge to a stable ratio *S*(*t*)/*D*(*t*) = const for *t* → ∞, given by:

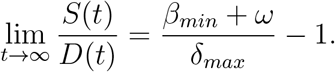

#### 4. Differences in relaxation dynamics of different model topologies after perturbations

In this section, we compare the relaxation dynamics of the four remaining model topologies after perturbations from steady-state, again using simple piecewise linear feedback functions to keep analyses tractable. We examine three different kinds of perturbations: first, removing differentiated cells; second, removing stem cells; and third, both of these perturbations combined. The first of these perturbations corresponds to a loss of differentiated cells which e.g. may arise as a consequence of physical stresses, inflammation or interactions of intestinal cells with microbes or parasites [9]. The second perturbation describes the loss of intestinal stem cells, which for instance arises as a consequence of radiation-induced cell damage [34]. Alternatively, stem cells numbers also might be reduced due to temporarily increased stem cell differentiation due to stochastic effects which may arise in such small populations.

Because of bilinear terms in the equations, we cannot in all cases obtain exact analytical solutions of the occurring dynamics, but use approximations of the dynamics based on a linearisation of the systems around their respective non-trivial steady-state (see Supplement for details).

The Jacobians and their Eigensystems for the remaining four models are shown in Figure 2, along with schematic model sketches and exemplary numerical simulations, illustrating typical solutions.

**FIG. 2.**
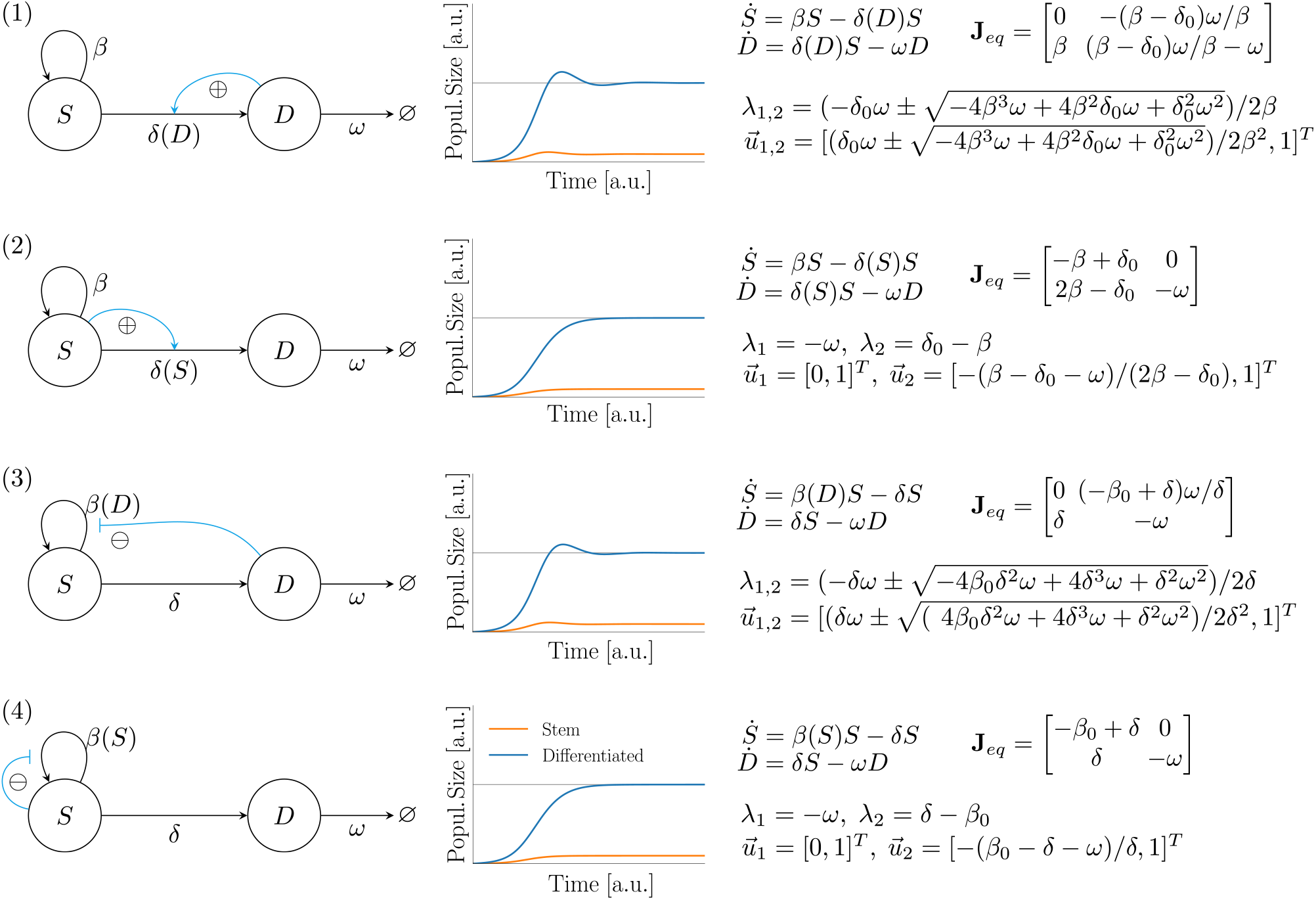
All four one-looped model topologies that can show homoeostasis. First column: schematic sketches; second column: exemplary numerical simulations; third column: model equations, as well as Jacobian at the non-trivial steady-state and its eigensystem for the case of a linear feedback function *β*(*x*) = *β*_0_ + *β_slope_x* or *δ*(*x*) = *δ*_0_ + *δ_slope_x*, respectively.

Physiologically, it is important that the number of differentiated cells recovers quickly, as these are responsible for carrying out the function of a tissue. We hence compare the models based on their ’defect’ χ of differentiated cells after perturbation – see Panel (a) of Figure 3 for an illustration. We define the defect as the total area between *D*(*t*)a nd 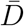 after a perturbation at *t* = 0, i.e.

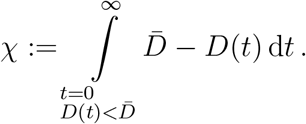

**FIG. 3.**
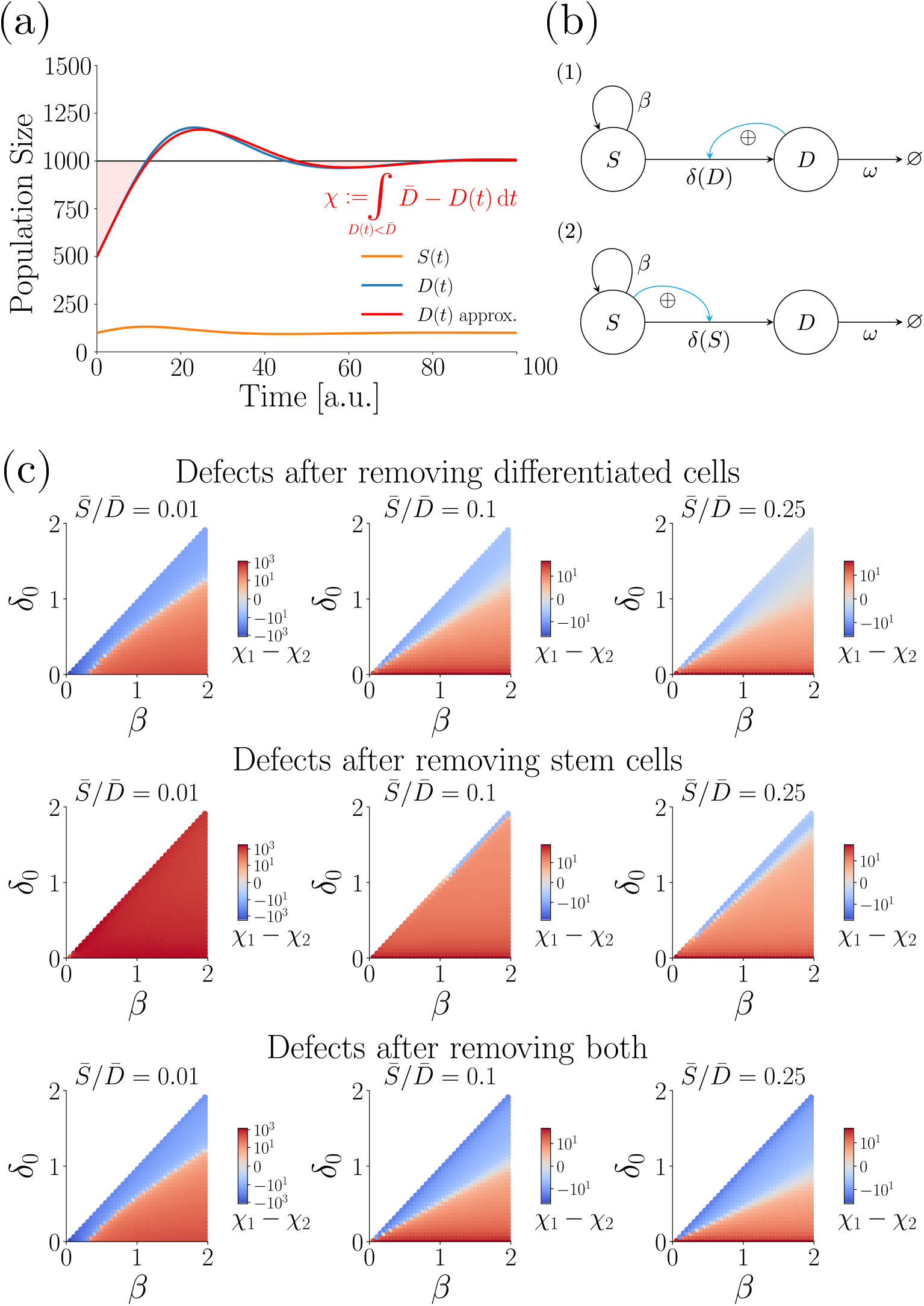
**(a)** Illustration of our model comparison: After a perturbation at *t* = 0, the model relaxes to its steady-state (continuous blue and orange lines, depicting differentiated and stem cells, respectively). We use a first-order approximation (continuous red line) to not have to rely on costly numerical solutions of the system, and compute the ’defects’ χ of the models we want to compare (shaded red area). **(b)** The two models we compare in this figure. Top: model 1, where stem cell differentiation is stimulated by the differentiated cell compartment; bottom: model 2, where the stem cell compartment stimulates its own differentiation. Note that model 1 is equivalent to our basic colon epithelium model derived earlier. **(c)** Difference in defects of model 2 and 1 throughout the parameter space. Areas shaded in red depict regions where model 1 shows a bigger defect, i.e. where the colon model recovers less gracefully than the alternative model; blue areas indicate the opposite. Columns represent different cases of stem cell fraction at steady-state (1, 10, and 25% respectively), rows represent the three different kinds of perturbations (removing differentiated cells, removing stem cells, and removing both, respectively).

The defect integrates only those time intervals in which 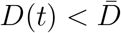, as this defines the lack of tissue functionality.

For all four models and the three types of perturbations, we derived either an analytical expression for the defect, or – if these are too complicated for a meaningful interpretation – an approximate solution (see Supplement). Three relations simplify their analysis: First, for all models we obtain 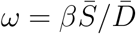. Second, the slope of the feedback function (*β_slope_* or *δ_slope_*, respectively) is not required for describing the dynamics relative to the steady-state, as the slope is not part of the Eigensystems of the steady-state Jacobians of the models. Third, the displacements of all models from steady-state of the first and second perturbation are linear in the initial displacement (Δ*D*(0) or Δ*S*(0), respectively). This indicates that in both cases the choice of initial displacement will not affect the comparison of the model defects, and we can express all defects as multiples of initial displacement. In case of the third perturbation, we remove the same fraction of stem and differentiated cells, and we express the defect as multiples of the intial displacement of differentiated cells. Overall, the complete parameter space we need to consider thus consists only of the ratio of stem to differentiated cells at steady-state and two additional free parameters (depending on the model *β*_0_, *δ* or *β*, *δ*_0_).

##### a. Colon model vs. direct stimulation of stem cell differentiation

We start by comparing our basic colon epithelium model (model 1), with model 2, where stem cell differentiation is stimulated by the stem cell compartment itself (Figure 3b). Figure 3c shows the difference in model defects throughout the parameter space (see Supplement for the raw values), where areas shaded in red indicate regions where the colon epithelium model has a bigger defect than the alternative model. In case of removing differentiated cells (first row) or removing both differentiated and stem cells at the same time (third row), there exist ample regions (shaded in blue) where the colon model recovers more efficiently – namely, whenever *δ*_0_ is close to *β*. This makes sense, because in the colon epithelium model removing differentiated cells will cause the stem cells to differentiate more slowly and thus grow in numbers quickly, thus being able to replenish the differentiated cell compartment more quickly. However, in case of removing only stem cells (second row), the colon epithelium model always performs worse than the alternative model, except for cases where the fraction of stem cells at the steady-state becomes sufficiently big (middle and right column). However, even then the difference between *δ*_0_ and *β* still needs to be small for the colon model to recover more efficiently.

##### b. Colon model vs. indirect inhibition of stem cell cycling rate

Next, we compare the recovery dynamics of the colon epithelium model with alternative model 3, where the differentiated cell compartment inhibits stem cell cycling. Because the two models have different system parameters (*β*, *δ*_0_ vs. *β*_0_, *δ*) we cannot compare them pointwise in parameter space. However, we can still compare the ranges of model defects for the three different perturbations and steady-state stem cell fractions (1%, 10%, and 25%) if we systematically vary the two other free parameters within biologically plausible intervals. In all cases, the defects of the two models fall into similar ranges (see Supplement). Hence, there does not seem to be any relevant difference in recovery behaviour between indirectly regulating stem cell differentiation vs. indirectly regulating stem cell cycling rate.

##### c. Colon model vs. self-inhibition of stem cell cycling rate

Finally, we compare the recovery dynamics of the colon epithelium model with the behaviour of model 4, where the stem cell compartment inhibits its own cycling rate. We again face the problem of the two models having some different system parameters (*β, δ*_0_ vs. *β*_0_, *δ*), thus we again compare the ranges of defects occurring in different scenarios. For the first and third perturbation, model 4 recovers slower by several orders of magnitude if its stem cell differentiation rate *δ* is small (see Supplement), since a small differentiation rate will cause the system to take longer to replenish the pool of differentiated cells. The only exception is when the steady-state stem cell fraction is very large. In contrast, if differentiated cells are removed from the colon epithelium model, the stem cell differentiation rate will decrease, causing the stem cell compartment to temporarily overshoot, until the growing differentiated stem cell compartment stimulates differentiation again. This way, the colon epithelium model is able to recover more quickly. For the remaining case of removing stem cells, the model defects fall into similar ranges. However the colon epithelium model shows its largest defect in case of a small *δ*_0_, whereas the alternative model performs worst in case of a very small difference between *β*_0_ and *δ*.

### C. Dedifferentiation improves recovery from perturbations, however offers an additional route of losing homoeostasis

Previously, we have seen that upon loss of stem cells, the colon epithelium model often does not recover as gracefully as model 2 (Figure 3), as removing stem cells will not alter the rate of stem cell differentiation. Hence, removing stem cells will lead to a significant loss of differentiated cells first, before differentiation rate drops enough for the stem cell compartment to replenish itself. This way, the transient behaviour is characterised by large oscillations in the number of differentiated cells, causing a large model defect.

We now study how allowing for the dedifferentiation of differentiated cells affects these recovery dynamics (Figure 4a). We again assume for simplicity that the rate of stem cell differentiation is given by a linear function with intercept *δ*_0_ and slope *δ_slope_*. We also assume that the function saturates to a maximum value *δ_max_* after some value of D. Hence, we have *δ*(*D*) = min{*δ*_0_ + *δ_slope_D*, *δ_max_*}, where *δ*_0_, *δ_slope_*, *δ_max_* > 0. Next, we also for now assume the same, but horizontally mirrored shape for *ϱ*, giving *ϱ*(*S*) = max{*ϱ*_0_ + *ϱ_slope_S*, *ϱ_min_*}, where *ϱ*_0_ > 0 *ϱ_slope_* < 0 *ϱ_min_* ≥ 0.

**FIG. 4.**
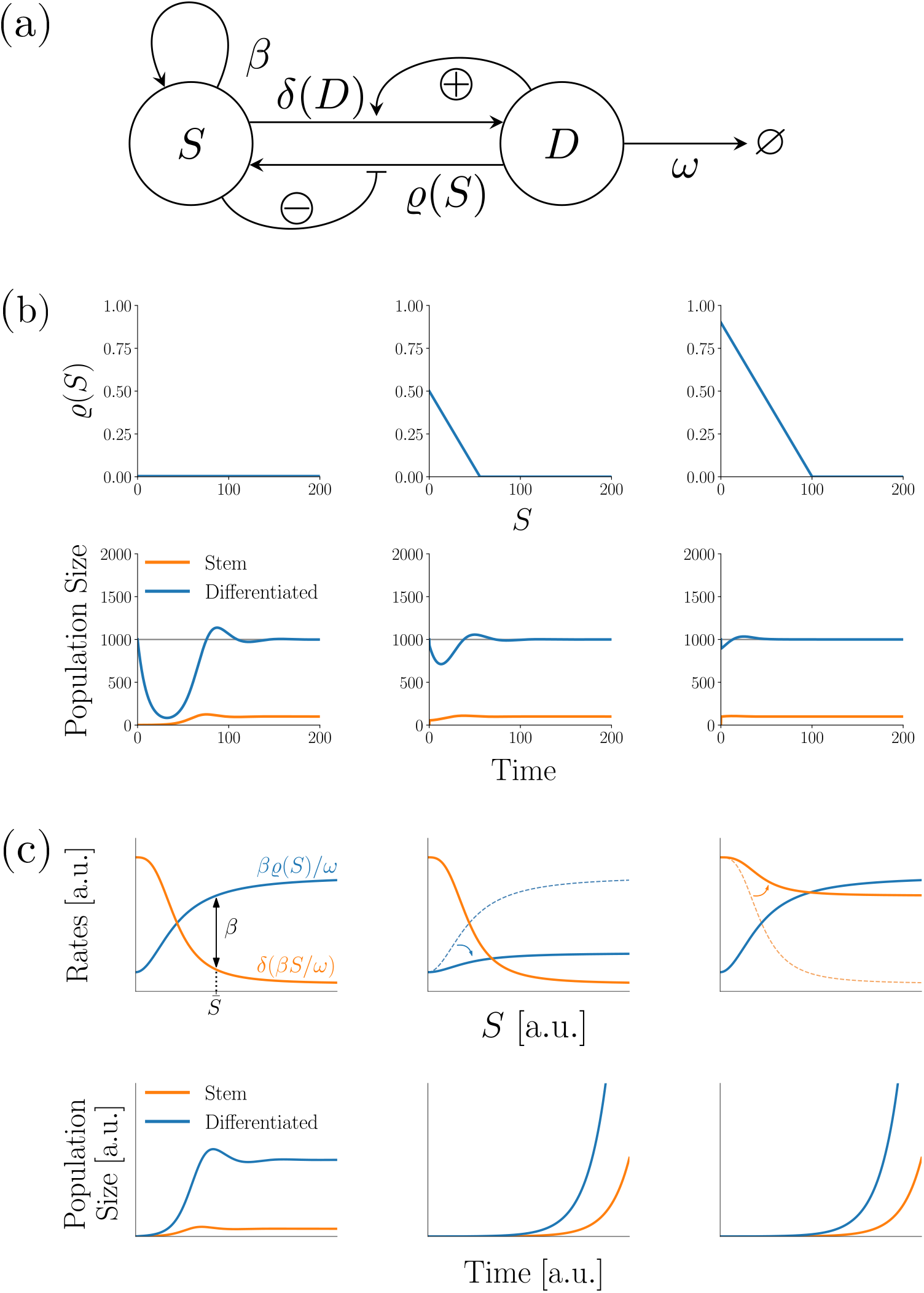
The colon epithelium model with dedifferentiation and its most important properties. **(a)**: Schematic sketch of the model. **(b)**: Exemplary numerical simulations of the system with piecewise linear differentiation rate function *δ* and different dedifferentiation rate functions *ϱ*: no dedifferentiation (first column), linear dedifferentiation with *ϱ*_0_ = 0.5, *ϱ_slope_* = −0.01 (second column) and a faster linear dedifferentiation *ϱ*_0_ = 0.9, *ϱ_slope_* = −0.01 (third column). System parameters are *β* = 1, *δ*_0_ = 0.9, *δ_slope_* = 10^−4^, *ω* = 0.1. Note how adding dedifferentiation, as well as increasing the higher maximum dedifferentiation rate *ϱ*_0_ makes the system recover more gracefully after removing stem cells. **(c)**: Adding dedifferentiation opens up a second way the system can lose homeostatic stability. First column shows the case of homoeostasis. Equilibrium stem cell pool size 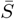 is given by solving 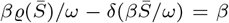. Second column: Sufficient decrease of differentiation rate destroys the non-trivial steady-state and unbounded growth occurs. Third column: Sufficient increase of dedifferentiation rate has the same consequence.

For brevity purposes, the exact calculations are presented in the Supplement. Briefly, adding a linear dedifferentiation function causes a faster decay of the oscillations after perturbations. Additionally, we can find a critical value 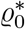, which, when exceeded by *ϱ*_0_ will also reduce the frequency of oscillations after perturbations. It is given by

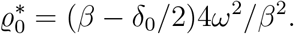

By means of a Taylor expansion around the steady-state, we can also generalise this finding to arbitrary decreasing differentiable functions *ϱ*. Figure 4b shows some exemplary numerical simulations of this behaviour. Observe, how the transient period after removing stem cells is characterised by smaller oscillation amplitudes and frequencies when dedifferentiation is allowed.

Next, we the influence of dedifferentiation on the stability of the non-trivial steady-state of the system. Regardless of the concrete functions *δ*, *ϱ*, we find that the non-trivial steady-state needs to satisfy 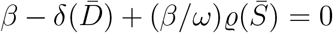 and 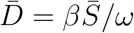. Hence, a non-trivial steady-state exists, if and only if we can solve

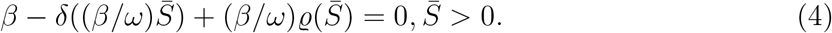

Importantly, this shows that dedifferentiation enables the system to lose its non-trivial steady-state through a different route, namely via a sufficient increase of the values of *ϱ* (see Panel (c) of Figure 4 for an illustration).

Then, the system will again converge to a stable cell type composition

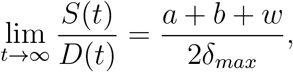

where 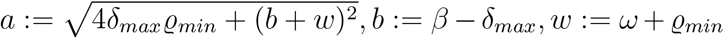 (see Supplement for details).

## IV. DISCUSSION

In this work, we derived and analysed a simple population dynamics model of the colon epithelium, taking into account the stimulating effect of differentiated cells onto the differentiation of stem cells [11–15]. Despite its simplicity, the model recapitulated known experimental results, such as the necessity to control the stem cell compartment for tissue homoeostasis [6], the balance between stem cell proliferation and differentiation required for constant tissue size [35, 36], and the emergence of stable intratumoural heterogeneity in a growing cancer (see below).

We showed that at any stable steady-state the number of differentiated cells is not affected by changes in their apoptosis rate, which is a biologically useful property, since differentiated cells are responsible for carrying out the primary function of a tissue [1, 10] and changes in apoptosis rate may regularly happen locally as a consequence of infections or mechanical wounding [9]. Additionally, we have seen that strong alterations in system parameters are required for the steady-state to be destroyed and unbounded growth to occur. In contrast, other single alterations will only change the position of the stable steady-state. This is equivalent to the case of a pre-cancerous lesion, where only a subset of ’canonical’ colorectal cancer mutations is yet present and which shows a higher number of cells, but requires further alterations to switch to unbounded growth [37]. Hence, despite its simplicity, our model recapitulates the observation of colorectal tumorigenesis being a characteristic multi-step process of subsequent system alterations in vivo [38], showcasing the intrinsic resilience of the system towards isolated mutations affecting system properties.

However, the indirect feedback loop in the model may cause oscillations. This is reminiscent of e.g. the dampened oscillations observed in the healthy haematopoietic system [39]. In case of very steep feedback functions, these oscillations are more pronounced or can even become undampened; thus, we may expect rather flat feedbacks in nature. Oscillations in cell numbers do not serve any immediately obvious biological purpose and indeed may not be desirable for upholding tissue homoeostasis. This led us to the question whether the colon model topology may offer additional beneficial properties compared to other topologies, to which we compared our model.

We revealed pronounced differences of the plausible model topologies regarding their recovery after perturbations. We have shown the existence of ample regions in parameter space where the colon epithelium model recovers more gracefully than the other examined model or at least equally well. If, however, a perturbation removes stem cells, recovery behaviour was poor.

This prompted us to study the behaviour of the model if we allow for dedifferentiation of differentiated cells. We revealed that this lets the model recover more gracefully; however, dedifferentiation may also cause the destruction of the non-trivial steady-state. This is biologically interesting because it suggests that a sufficient degree of dedifferentiation may offer an alternative route towards unbounded growth. Experimental data seems to confirm the possibility of tumours arising from the differentiated intestinal epithelium [40]. Interestingly, the mutations required for this indeed occur rather often in colorectal cancers [41, 42], suggesting that increased dedifferentiation might regularly contribute to tumorigenesis. Experimental data [43] point into the same direction.

Obviously, dedifferentiation interferes with the invariance of the steady-state number of differentiated cells to changes in apoptosis rate. However, if the dedifferentiation rate around the steady-state is small and only becomes noticeable if a sizeable fraction of stem cells is removed (which seems to be suggested by the literature, see Tata et al. [3] and Beumer and Clevers [8]), then around the steady-state dedifferentiation becomes negligible and the biologically desirable invariance still holds.

We also revealed that a tissue can be expected to converge to a stable cell-type ratio during unbounded growth. From a biological point of view, this relates to the topic of intra-tumoural heterogeneity [44–46], and suggests that in case of cancerous growth the colon epithelium can still be expected to recapitulate known cellular hierarchies and differentiation gradients. In case of breast cancer [47] and cancers of the haematopoietic system [48] this has already been observed experimentally.

An interesting avenue for future research lies in comparing healthy with tumoural intestinal tissue to clarify, whether one indeed observes similar cellular subpopulations and differentiation gradients. Recent research indeed suggests this [49, 50]. Tracking the numbers of different cell types over time during development and after perturbations could therefore be expected to yield data for validating and parametrising the presented model. Additionally, comparing the quantitative behaviour of tissues in different stages of tumorigenesis could elucidate at which stage which system properties change in which way, and thus increase our mechanistic insight into the dynamics of tumour development. This, in turn, might ultimately help to contribute to the development of novel approaches to limiting tumoural growth *in vivo*.

## ACKNOWLEDGMENTS

We thank Christine Sers and Markus Morkel for fruitful discussions and helpful comments during an early stage of this project and DFG for funding (RTG2424, CompCancer). MMF would like to thank J. F. W. Maas for help during the preparation of the schematic model sketches.

## DECLARATIONS

### A. Author contributions

All authors conceived of the presented ideas. MMF carried out model derivations and analyses, and wrote the initial version of the manuscript, both with help from NB and HH. HH and NB contributed to the final version of the manuscript. NB supervised the project. All authors have read and approve of the final version of this manuscript.

