## Supplementary material for "Maintaining and escaping feedback control in hierarchically organised tissue: a case study of the intestinal epithelium": Supplmentary Materials

### I. MODEL DYNAMICS AFTER PERTURBATIONS

Let $(\Delta S(0), \Delta D(0))$ denote a perturbation away from the steady-state $(\bar{S}, \bar{D})$ of a system at $t = 0$. Let $\vec{q}(t) = [\Delta S(t), \Delta D(t)]^T$ denote the dynamics of the displacement after the perturbation. We can approximate to first order $\mathrm{d}\vec{q}(t)/\mathrm{d}t \approx \mathbf{J}_{eq}\vec{q}(t)$, where $\mathbf{J}_{eq}$ denotes the Jacobian of the system at the steady-state. This linear system is then solved by $\vec{q}(t) = c_1 e^{\lambda_1 t}\vec{u}_1 + c_2 e^{\lambda_2 t}\vec{u}_2$, where $\lambda_{1,2}$ denote the eigenvalues of $\mathbf{J}_{eq}$, $\vec{u}_{1,2}$ denote their corresponding eigenvectors, and $c_{1,2} \in \mathbb{C}$ are to be chosen to satisfy the initial condition $\vec{q}(0) = [\bar{S} + \Delta S(0), \bar{D} + \Delta D(0)]^T$.

Because model 1 may show oscillations during its relaxation, an analytical calculation of its defects is theoretically possible, but leads to expressions too complicated to handle and meaningfully interpret. For this reason, we instead only derive the approximative relaxation dynamics of the model for the three perturbations and will calculate the corresponding model defects numerically. We get:

$$\Delta D_1(t) = \begin{cases} \Delta D(0)\dfrac{\cosh(rt) - (d/r)\sinh(rt)}{e^{dt}} & \text{First perturbation} \\ \Delta S(0)\beta\dfrac{\sinh(rt)}{re^{dt}} & \text{Second perturbation} \\ \Delta D(0)\dfrac{\cosh(rt) - (d/r)\sinh(rt)}{e^{dt}} + \Delta S(0)\beta\dfrac{\sinh(rt)}{re^{dt}} & \text{Third perturbation,} \end{cases}$$

where $d := \delta_0\omega/(2\beta)$, and $r := \sqrt{-4\beta^3\omega + 4\beta^2\delta_0\omega + \delta_0^2\omega^2}/(2\beta)$.

Next, we calculate the defects $\chi_2$ of model 2 for the three perturbations. For the case of the first perturbation, the defect can be directly obtained by solving the system analytically for $S(0) = \bar{S}, D(0) = 0$ and calculating the integral. The other two defects are obtained by using the first-order approximation of the dynamics derived earlier and analytically calculating their integrals after choosing the respective initial conditions. We get:

$$\chi_2 = \begin{cases} \Delta D(0)\dfrac{1}{\omega} & \text{First perturbation} \\ \Delta S(0)\dfrac{2\beta - \delta_0}{\beta - \delta_0 - \omega}\left(1/(\delta_0 - \beta) + 1/\omega\right) & \text{Second perturbation} \\ \Delta D(0)\dfrac{1}{\omega} + \Delta S(0)\dfrac{2\beta - \delta_0}{\beta - \delta_0 - \omega}\left(1/(\delta_0 - \beta) + 1/\omega\right) & \text{Third perturbation.} \end{cases}$$

For model 3, we again only derive the relaxation dynamics after the three perturbations, and will examine them numerically. They are given by:

$$\Delta D_3(t) = \begin{cases} \Delta D(0)\dfrac{\cosh(rt) - (d/r)\sinh(rt)}{e^{dt}} & \text{First perturbation} \\ \Delta S(0)\dfrac{\delta \sinh(rt)}{re^{dt}} & \text{Second perturbation} \\ \Delta D(0)\dfrac{\cosh(rt) - (d/r)\sinh(rt)}{e^{dt}} + \Delta S(0)\dfrac{\delta \sinh(rt)}{re^{dt}} & \text{Third perturbation,} \end{cases}$$

where $d := \omega/2$, and $r := \sqrt{\omega(-4\beta_0 + 4\delta + \omega)}/2$.

For model 4, we can analytically calculate the defects after the three perturbations. They are:

$$\chi_4 = \begin{cases} \Delta D(0)\dfrac{1}{\omega} & \text{First perturbation} \\ \Delta S(0)\dfrac{\delta}{(\beta_0 - \delta)\omega} & \text{Second perturbation} \\ \Delta D(0)\dfrac{1}{\omega} + \Delta S(0)\dfrac{\delta}{(\beta_0 - \delta)\omega} & \text{Third perturbation.} \end{cases}$$

#### II. MODEL DEFECTS AFTER PERTURBATIONS

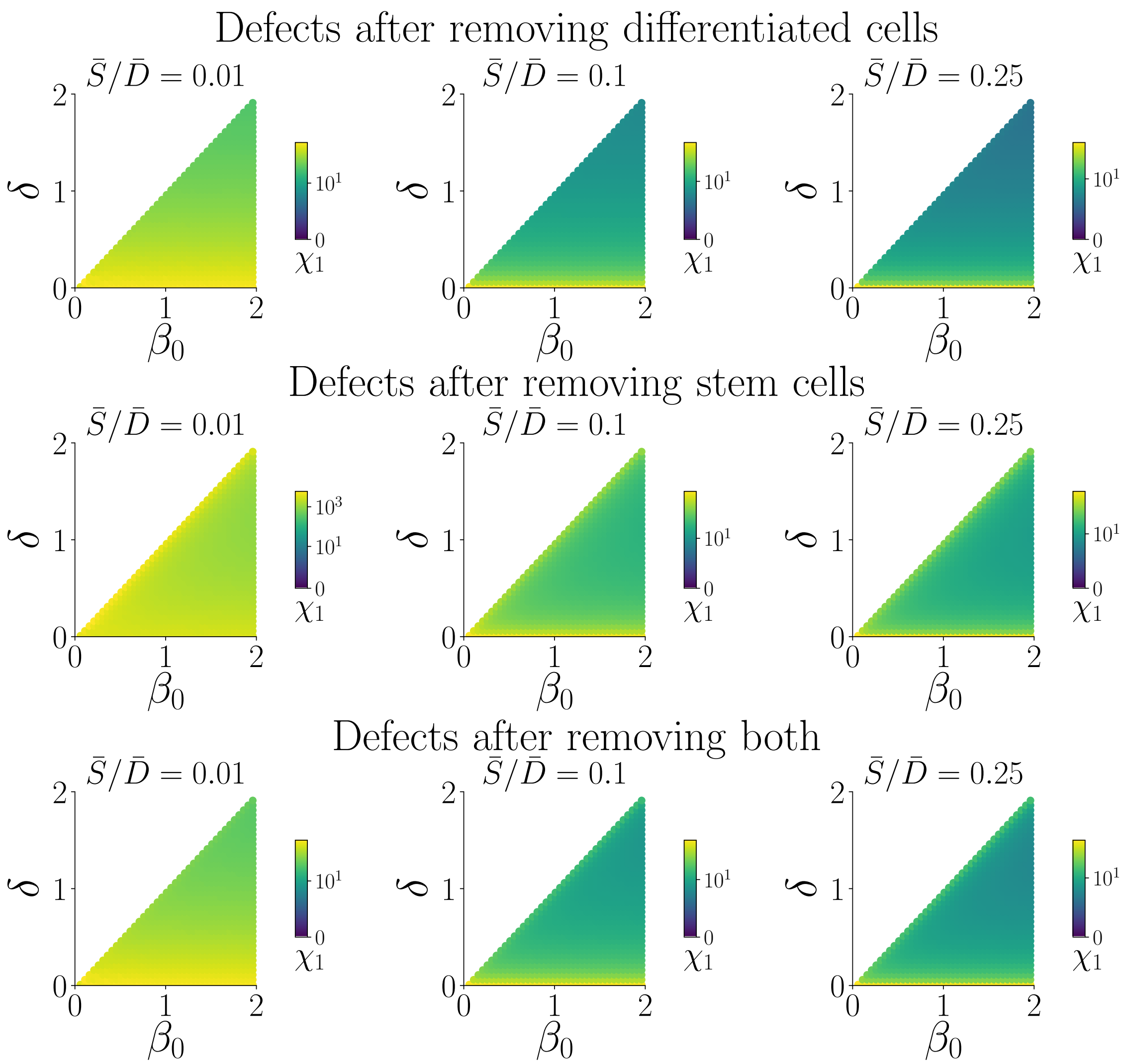


FIG. 1. The model defects $\chi_1$ of model 1, the colon epithelium model, throughout its parameter space. Defects are given in multiples of initial perturbation size. Columns represent three different scenarios with different steady-state stem cell fractions of 1%, 10%, and 25%, respectively.

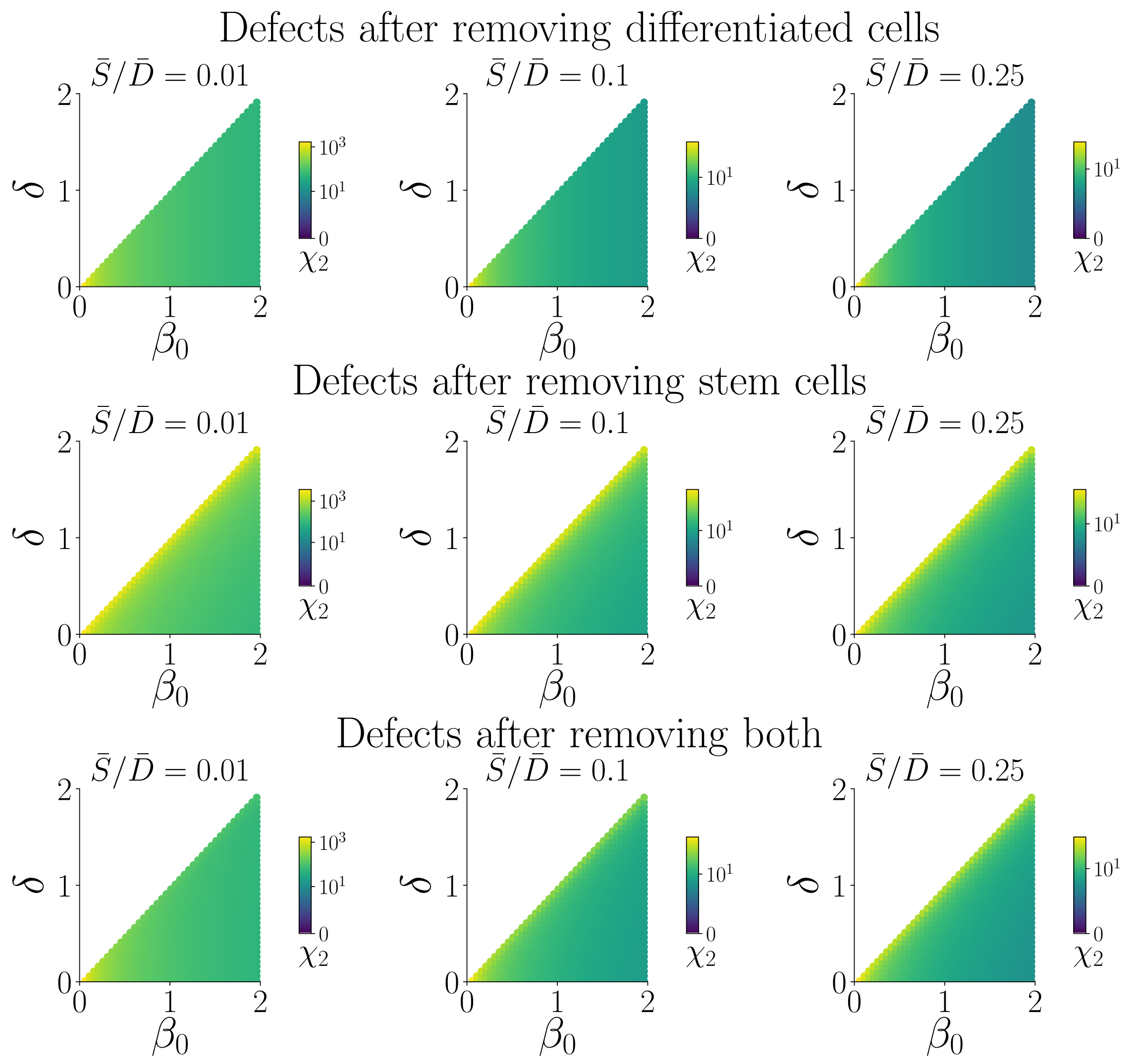


FIG. 2. The model defects $\chi_2$ of model 1, the colon epithelium model, throughout its parameter space. Defects are given in multiples of initial perturbation size. Columns represent three different scenarios with different steady-state stem cell fractions of 1%, 10%, and 25%, respectively.

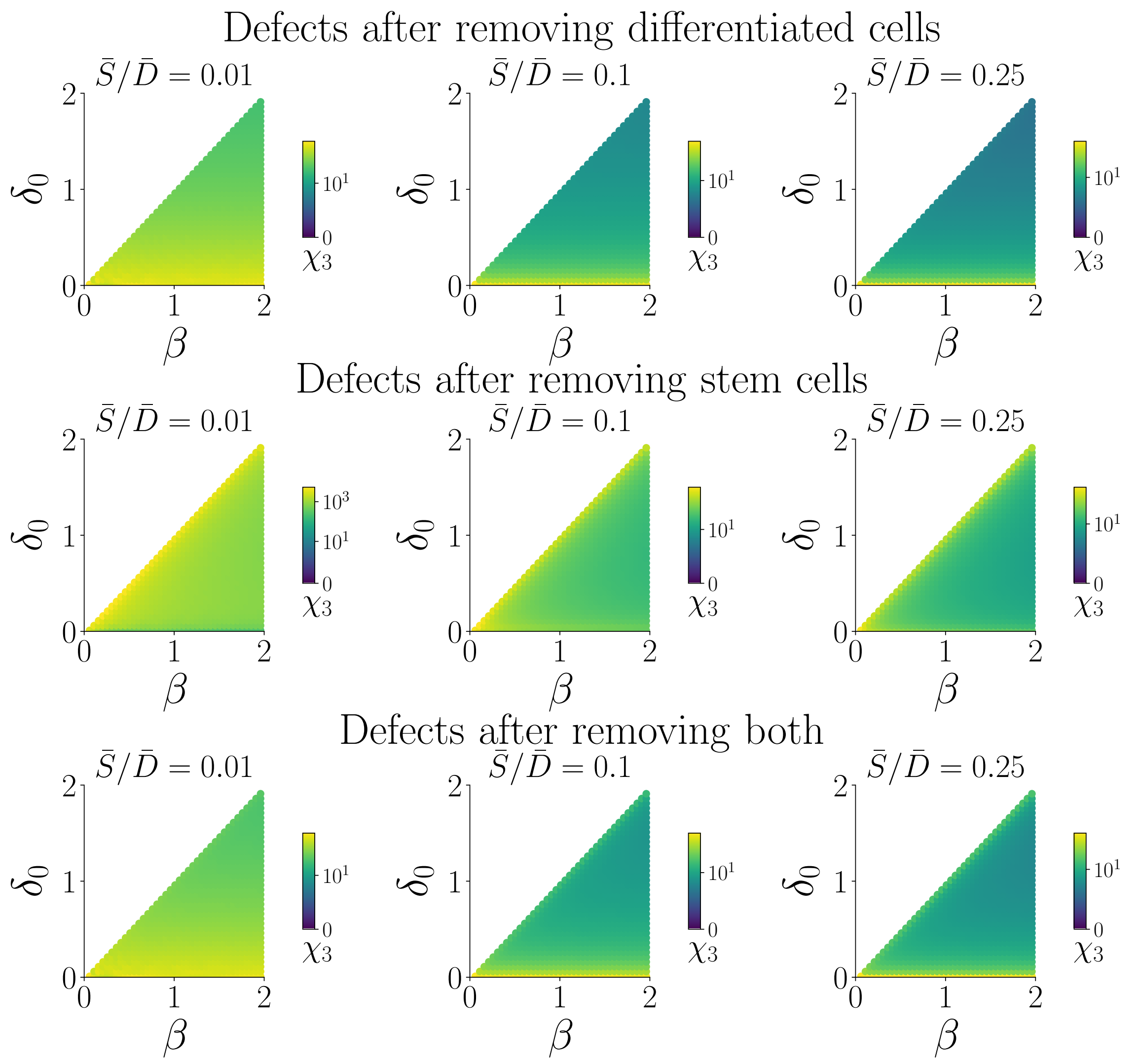


FIG. 3. The model defects $\chi_3$ of model 3, where stem cell cycling rate is controlled by the number of differentiated cells, throughout its parameter space. Defects are given in multiples of initial perturbation size. Columns represent three different scenarios with different steady-state stem cell fractions of 1%, 10%, and 25%, respectively.

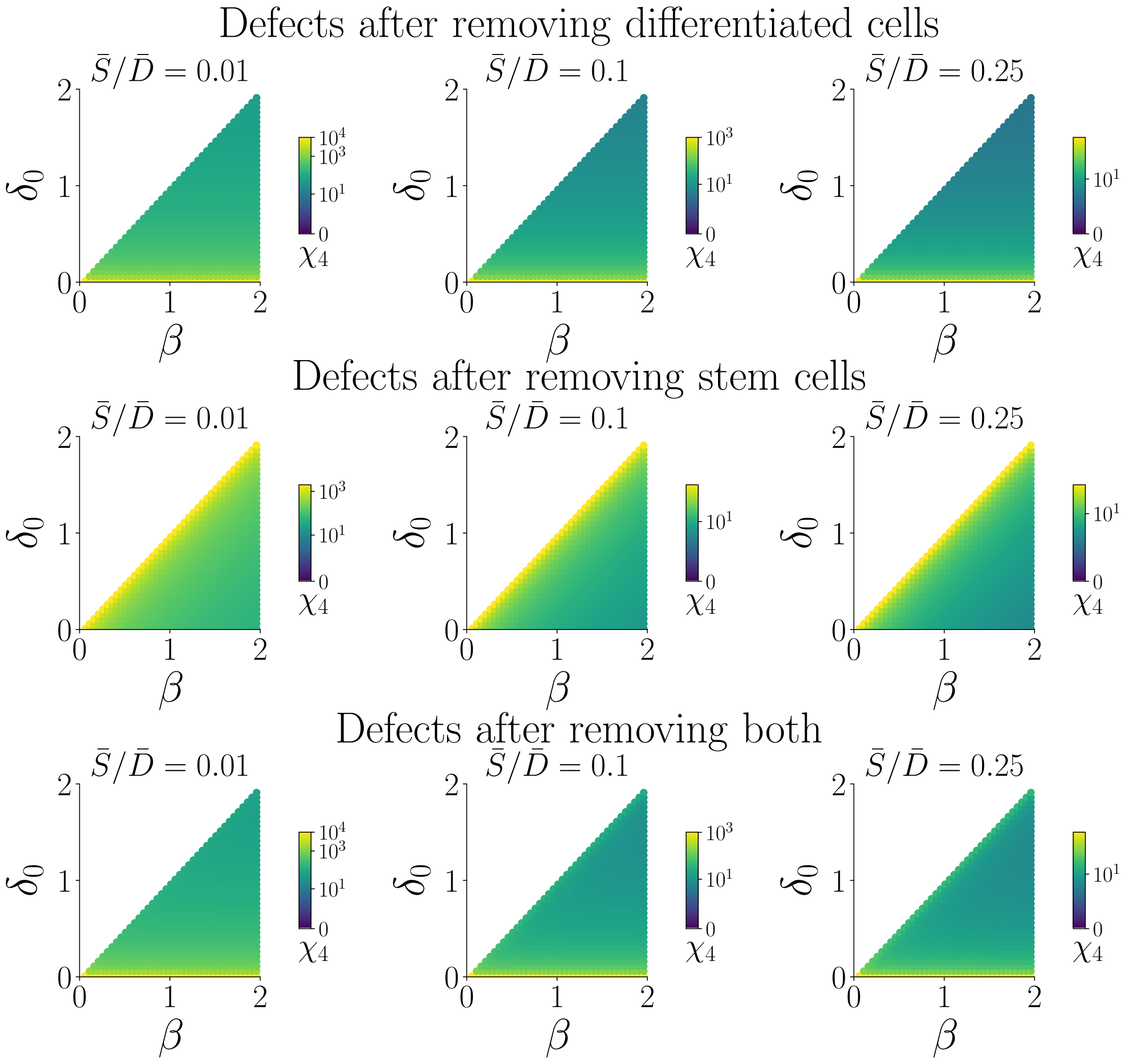


FIG. 4. The model defects $\chi_4$ of model 4, where the stem cell compartment inhibits its own cycling rate, throughout its parameter space. Defects are given in multiples of initial perturbation size. Columns represent three different scenarios with different steady-state stem cell fractions of 1%, 10%, and 25%, respectively.

#### III. DEDIFFERENTIATION CAN LIMIT OSCILLATORY BEHAVIOUR

The modified model permits exactly one non-trivial steady-state at

$$\bar{S} = -\frac{(\beta - \delta_0)\omega + \varrho_0 \beta}{\beta(\varrho_{slope} - \delta_{slope})};\ \bar{D} = \beta \bar{S}/\omega,$$

and its Jacobian at this steady-state is given by

$$\mathbf{J}_{eq} = \begin{pmatrix} -\frac{\varrho_0 \beta}{\omega} & -\frac{(\beta - \delta_0)\omega}{\beta} \\ \beta + -\frac{\varrho_0 \beta}{\omega} & \frac{(\beta - \delta_0)\omega}{\beta} - \omega. \end{pmatrix}$$

This matrix has eigenvalues

$$\lambda_{1,2} = \frac{-\beta^2 \varrho_0 - \delta_0 \omega^2 \pm \sqrt{(\beta^2 - \varrho_0 + \delta_0 \omega^2)^2 - 4(\beta^3 \varrho_0 \omega^2 + \beta^3 \omega^3 - \beta^2 \delta_0 \omega^3)}}{2\beta\omega}$$

If oscillations occur, we have complex eigenvalues, which we can split into a real and an imaginary part as follows:

$$\Re(\lambda_{1,2}) = -\frac{\beta \varrho_0}{2\omega} - \frac{\delta_0 \omega}{2\beta};\ \Im(\lambda_{1,2}) = \frac{\sqrt{(-\beta^2 - \varrho_0 + \delta_0 \omega^2)^2 + 4(\beta^3 \varrho_0 \omega^2 + \beta^3 \omega^3 - \beta^2 \delta_0 \omega^3)}}{2\beta\omega}.$$

For the colon epithelium model without dedifferentiation the real part of the complex eigenvalues of the Jacobian at steady-state was given by $-\delta_0\omega/(2\beta)$. Accordingly, additionally allowing for dedifferentiation reduces the real part by $\beta\varrho_0/(2\omega) > 0$, hence always causing a faster decay of the oscillations after perturbations.

For the model without dedifferentiation we had an imaginary part of the eigenvalues of $\sqrt{4\beta^3\omega - 4\beta^2\delta_0\omega - \delta_0^2\omega^2}/(2\beta)$. Hence, adding the dedifferentiation to the model changes the radicand of the imaginary part of the eigenvalues by

$$\Delta = \beta \varrho_0 - \frac{\beta^2 \varrho^2}{4\omega^2} - \frac{\varrho_0 \delta_0}{2}.$$

Thus, we can find a critical value $\varrho_0^*$, which, when exceeded by $\varrho_0$ will reduce the frequency of oscillations after perturbations. It is given by

$$\varrho_0^* = (\beta - \delta_0/2)4\omega^2/\beta^2.$$

In other words, adding a linear dedifferentiation function will speed up the amplitude decay of the oscillations after perturbations and can also, in case of a sufficiently big basal dedifferentiation rate, decrease the frequency of these oscillations.

We can generalise this finding to arbitrary decreasing differentiable functions $\varrho$. To this end, we construct a Taylor expansion of $\varrho$ around the steady-state of the form $\varrho(D) \approx a + bD + \mathcal{O}(D^2)$ with $a, b \in \mathbb{R}$. This way, in a sufficiently small neighbourhood around the steady-state, the system behaves as if $\varrho$ was linear; and because $\varrho$ by definition is always positive and monotonically decreasing, clearly $b < 0$ and accordingly $a > 0$. Hence, the argument for linear functions $\varrho$ we made previously also applies here when we simply replace $\varrho_0$ with $a$.

#### IV. CONVERGENCE OF THE COLON EPITHELIUM MODEL WITH DEDIFFERENTIATION TO A STABLE CELL TYPE RATIO

For sufficiently high population sizes, the dynamics of the system are governed by the set of linear differential equations

$$\frac{\mathrm{d}S(t)}{\mathrm{d}t} = \beta S(t) - \delta_{max} S(t) + \varrho_{min} D(t)$$
$$\frac{\mathrm{d}D(t)}{\mathrm{d}t} = \delta_{max} S(t) - \varrho_{min} D(t) - \omega D(t).$$

For convenience, we define $b := \beta - \delta_{max}$, $w := \omega + \varrho_{min}$, $r := \varrho_{min}$, $d := \delta_{max}$, giving

$$\frac{\mathrm{d}S(t)}{\mathrm{d}t} = bS(t) + rD(t)$$
$$\frac{\mathrm{d}D(t)}{\mathrm{d}t} = dS(t) - wD(t).$$

This system has the general solution

$$S(t) = \frac{(a-b-w)S_0 - 2rD_0 + ((a+b+w)S_0 + 2rD_0)e^{at}}{2ae^{\frac{a-b+w}{2}t}}$$

$$D(t) = \frac{-2dS_0 + (a+b+w)D_0 + (2dS_0 + (a-b-w)D_0)e^{at}}{2ae^{\frac{a-b+w}{2}t}},$$

where $a := \sqrt{4dr + (b+w)^2}$. Taking the limit yields

$$\lim_{t\to\infty} \frac{S(t)}{D(t)} = \frac{a+b+w}{2d}.$$
